# Detecting GPCR Complexes in Postmortem Human Brain with Proximity Ligation Assay and A Bayesian Classifier

**DOI:** 10.1101/687780

**Authors:** Ying Zhu, József Mészáros, Roman Walle, Rongxi Fan, Ziyi Sun, Andrew J. Dwork, Pierre Trifilieff, Jonathan A. Javitch

## Abstract

Despite the general controversy regarding the existence and physiological relevance of Class A GPCR dimerization, there is substantial evidence for functional interactions between dopamine D2 receptor (D2R) and adenosine A2A receptor (A2AR). A2AR-D2R complexes have been detected in rodent brains by proximity ligation assay (PLA), but their existence in the human brain is yet to be demonstrated. In this study, we used brightfield PLA, combined with a systematic sampling and a parameter-free naïve Bayesian classifier, and demonstrated proximity between D2R and A2AR in the adult human ventral striatum, consistent with their colocalization within complexes and the possible existence of D2R-A2AR heteromers. These methods are applicable to the quantitative analysis of proximity of two proteins and the expression of individual proteins.

**Method Summary:** Brightfield proximity ligation assay (PLA) was used to assess the expression of G protein-coupled receptors and their proximity in postmortem adult human brains. A novel automated machine learning method (Bayesian Optimized PLA Signal Sorting) was developed to automatically quantify brightfield PLA data.

## Introduction

The dopamine receptor type 2 (D2R) is an extensively studied Class A G protein-coupled receptor (GPCR) that has been shown to play a critical role in various brain functions. Alterations in D2R signaling/activity have been described in a variety of neuropsychiatric disorders, such as schizophrenia [1, 2], Parkinson’s [3, 4], Alzheimer’s [5], Huntington’s diseases [7] and addiction [8]. Moreover, the D2R is the common target of all current antipsychotic medications [9], which has led to an extensive literature regarding its pharmacological and signaling properties [10]. The transduction signals downstream of D2R include Gα_i/o/z_βγ-mediated pathways, which inhibit adenylyl cyclase and modulate voltage-gated K^+^ and Ca^2+^ channels, as well as arrestin2/3-mediated pathways, which promote D2R internalization, terminate G protein signaling, and activate G protein independent signaling [11]. Previous studies in cell lines suggest that D2Rs might function as part of heteromeric complexes, within which D2R activity and signaling can be modulated by another receptor subunit [12-15]. The putative heteromer formed by the D2R with the adenosine A2A receptor (A2AR) is one of the most studied among Class A GPCRs [16-20], and the allosteric modulation observed in the D2R-A2AR complex has led to the hypothesis that pharmacological targeting of A2AR could be an efficient strategy to modulate D2R activity in pathological conditions, neurodegenerative disorders in particular [21, 22]. However, the structural properties of Class A receptor heteromers, their existence *in vivo*, and their relevance to receptor physiology or pathophysiology remain unclear and a topic of active study and debate [23-25]. The signaling properties of putative D2R-A2AR heteromers have been mostly studied in heterologous systems, and the existence of such receptor complexes in native mammalian brain is still poorly characterized.

Both time-resolved fluorescence resonance energy transfer (FRET) based assays [26, 27] and antibody-based *in situ* proximity ligation assays (PLA) [28, 29] have been used to study receptor complexes in native tissue. Using fluorescent PLA, we and others have successfully detected endogenous D2R-A2AR complexes in the rodent striatum, which provided *ex vivo* evidence for the existence of D2R-A2AR heteromeric complexes composed of native receptors [29, 30]. However, the existence of D2R-A2A complexes in human brains has yet to be established.

PLA has been widely used to assess protein-protein interaction, protein expression, and post-translational modification both *in vitro* and *ex vivo* [31-34]. PLA puncta are generated when a pair of oligonucleotide-conjugated antibodies bind to neighboring antigens, followed by ligation of the oligonucleotides and subsequent rolling cycle amplification, leading to DNA structure that can be detected by fluorophore- or horseradish peroxidase (HRP)-labeled oligonucleotide probes. In contrast to standard immunohistochemistry and immunofluorescence, both of which rely on a field of precipitate or fluorescence that can only be quantified by total intensity, PLA results in individual puncta that allow relative quantification of proteins (single PLA) or complexes (dual PLA) with higher spatial resolution. To date, PLA has been applied to postmortem human brain in a limited number of studies. PLA was used to assess alpha-synuclein oligomers in Parkinson’s disease [35] and to detect the interaction between sortilin-related receptor 1 (SORL1) and amyloid beta precursor protein (APP) in Alzheimer disease [36]. A limiting factor in applying these approaches to human brain relates to the size of the samples, which makes it challenging to obtain representative data while minimizing the number of samples, as classical stereology techniques are extremely time-consuming. Furthermore, PLA is highly sensitive to tissue processing [29], and its use in postmortem human brain tissue requires additional consideration and optimization. Notably, human brains are classically fixed by immersion in fixative and paraffin-embedded, which is not the method of choice for PLA in rodent brain tissue, and therefore has not been systemically explored.

Herein, we optimized *in situ* PLA to detect D2R, A2AR and D2R-A2AR complexes with brightfield microscopy in the human ventral striatum and developed a new approach combining whole-slide scanning, systematic random sampling and parameter-free automated image analysis that employs a naïve Bayesian classifier for faithful and robust signal separation. This study constitutes a proof-of-concept for systematic quantification of expression of individual proteins – as well as antigen complexes - from postmortem human brain samples.

## Materials and Methods

### Human Subjects

The study was approved by the Institutional Review Board (IRB) of New York State Psychiatric Institute (NYSPI, Protocol 6477R). Human brain specimens (Table 1) were from autopsies at the Institute for Forensic Medicine, Skopje, Macedonia. The cerebral hemispheres were sliced coronally at 2 cm intervals. Slices from the left hemisphere were placed in home-made cassettes and fixed in 10% formalin in phosphate buffer (PB, pH 7.4) at room temperature for 5 days, then rinsed in tap water and transferred to phosphate buffered saline (PBS) with 0.02% sodium azide at 4°C. The fixed slices were examined by a neuropathologist who dissected a standard series of blocks, which were dehydrated and infiltrated with paraffin in a vacuum infiltration processor [37]. Paraffin-embedded blocks of the rostral left striatum, including ventral putamen (Ptm), ventral caudate (Cdt), and nucleus accumbens (NAcc), were used in this study. Sections were cut at a thickness of 6 μm at the Core Facility, Department of Pathology, the Herbert Irving Comprehensive Cancer Center (HICCC), Columbia University Medical Center.

**Table 1.** Human Subject information

| Accession number | Days in PB buffer | SEX | Age | Cause of death | Postmortem Interval (hours from death to autopsy) | Blood Alcohol | Psychiatric Diagnosis | Comment | Brain Toxicology at CU | Neuropathology |
| --- | --- | --- | --- | --- | --- | --- | --- | --- | --- | --- |
| PI12260 | 1951 | F | 68 | Myocardial infarction, acute. | 13 | 0 | Major Depressive Disorder. Single Episode Chronic, Melancholic, with Moderate Severity | No history of medication for depression | Cotinine | Encephalomalacia, subacute, small, in left ventrolateral thalamus. Atherosclerosis, moderate. |
| PI12270 | 920 | M | 47 | Myocardial infarction, acute. | 19 | 0 | NONE |  | Cotinine | Hypertensive vasculopathy |
| PI12277 | 921 | M | 50 | Hemorrhage from internal and external injuries. Pedestrian struck by car | 21 | 0 | None | No frozen tissue | Not available | Small, acute hemorrhage in frontal horn of left lateral ventricle |

### Specimen preparation from mouse brains

All mice were handled in accordance with the National Institute of Health (NIH) Guide for the Care and Use of Laboratory Animals. Experimental protocol (NYSPI1388) was approved by the Institutional Animal Care and Use Committee (IACUC) at NYSPI. Whole brains were collected from adult mice (3 months old). The mice were deeply anesthetized with ketamine (100 mg/kg)/xylazine (10 mg/kg) mix for all the following tissue collection. Transcardial perfusion was performed with 4% paraformaldehyde (PFA) in PBS (pH 7.4). Brains were removed and post-fixed in 4% PFA overnight at 4 °C. Coronal sections (30 μm) of unembedded fixed tissue were generated with a vibrating blade on a LeicaVT 1200 slicer and stored in PBS with 0.02% sodium azide for up to one week until PLA. When appropriate, mouse brains were collected without transcardial perfusion, and fixed in 4% PFA for 4 days at 4 °C. The fixed brains were trimmed, dehydrated, embedded in paraffin, and sectioned (6 μm) at the Core Facility, Department of Pathology, HICCC, Columbia University Medical Center. To collect fresh frozen samples, brains were removed and snap frozen in chilled 2-methylbutane (−20°C), and stored at −80°C [38]. Frozen sections (20 μm) were prepared with a Cryostat (Leica CM3050S), mounted on positively charged glass slides (Fisher Scientific), air-dried, and fixed with 4% PFA in PBS for 10 mins before PLA.

### PLA on Brain sections

PLA on sections from mouse brains fixed by transcardial perfusion was performed in 24-well plates according to the published protocol for free-floating sections [29]. PLA on snap-frozen and paraffin-embedded brain tissue was performed on sections mounted on positively charged slides (Fisher Scientific). The paraffin sections for immunohistochemistry (IHC) and PLA were deparaffinized in xylene, rehydrated in serial denatured ethanol baths (Fisherbrand™, HistoPrep™), and rinsed twice briefly in 0.1M Tris-buffer with 0.9% saline (TBS). Antigen retrieval was performed by boiling brain sections in 10 mM sodium citrate buffer (pH 6.5) for 6 min (with a 5-min-interval in the microwave after the first 3 min and addition of buffer to compensate for evaporation). The sections for PLA brightfield (PLA-BF) were quenched with 1% H_2_O_2_ for 30 min to inactivate endogenous peroxidase. After brief rinses with TBS containing 0.1% Triton X-100 (TBS-T), sections were incubated with blocking buffer (Duolink blocking buffer for PLA) at room temperature for 1 hour, and then with primary antibodies diluted in the Duolink antibody dilution buffer at 4 °C overnight. Immunofluorescent PLA (PLA-FL) (red) and brightfield PLA (PLA-BF) were performed with the Duolink PLA Fluorescence protocol and Duolink BrightField protocol (Sigma) according to the manufacturer’s manual. In this study we performed both single-recognition PLA (single PLA), which allows detection of a single antigen with only one primary antibody, and dual-recognition PLA (dual PLA), which detects the D2R-A2AR complex with two primary antibodies.

Luxol fast blue and cresyl violet (LFB/CV) staining of the human serial paraffin sections were performed according to standard protocol at the HICCC to outline the sub-territories of the striatum.

### Antibodies

Anti-D2R (0.5mg/ml, rabbit polyclonal, Millipore, ABN462) was used at 1/300 for single PLA (recognition of one antigen), and 1/100 for dual PLA (proximity between 2 antigens). Anti-A2AR (0.5mg/ml, mouse monoclonal, Millipore 05-717) was used at 1/500 for single PLA, and 1/300 for dual PLA.

### Microscopy and Sampling

For fluorescent staining, images were taken with a Zeiss LSC510 confocal laser-scanning microscope, using a 63X oil objective and z-stack scanning with a step interval of 0.5 μm. Brightfield whole slide images were taken with a Leica SCN400 at HICCC, using tube lens magnification (2x) in addition to 20x objective lens (Numerical Aperture to achieve 40x magnification and z-stack scanning with a step interval of 1 μm. Whole-slide scanned (WSC) virtual images were viewed with Leica SCN400 image viewer (version 2.2) and a single layer with the most PLA puncta in focus was used to export images (zoom at 40x) for PLA signal quantification.

A systematic random sampling (SRS) method adapted from Gundersen et al [39] was performed to choose areas in the brain region of interest (ROI) to quantify PLA signal. Briefly, a grid was used to divide the ROI in virtual images into a series of sampling areas. Then one counting area (0.11 mm^2^, covering about 4% of the sampling area, scale 4000 pixels/mm) per sampling area, at the same location inside each sampling area, was exported for quantification (Table 2, Suppl. Fig. 1 and Suppl. Method). Therefore, all the counting areas are distributed evenly within the ROI, but the specific locations of the counting areas are randomly selected. All counting areas in the ROI were exported as RGB tiff files with the ROI export function in ScnViewer 2.2.

**Table 2.** Fraction of images to ROI

| Sample# | 0.4x scale<br>pixels/mm<sup>2</sup> | ROI (NAcc)<br>area (mm<sup>2</sup>) | Sampling<br>Area (mm<sup>2</sup>) | Counting area<br>(mm<sup>2</sup>) | Counting/<br>Sampling (%) | Number of Counting<br>images |  |  | Fraction of ROI (%) |  |  |
| --- | --- | --- | --- | --- | --- | --- | --- | --- | --- | --- | --- |
|  |  |  |  |  |  | Dual | A2A | D2R | Dual | A2A | D2R |
| PI12260 | 40.02 | 36.32 | 2.72 | 0.11 | 4.08 | 13 | 13 | 6 | 3.94 | 1.82 | 3.33 |
| PI12270 | 41.96 | 84.9 | 2.47 | 0.11 | 4.32 | 25 | 25 | 31 | 3.24 | 4.02 | 3.50 |
| PI12277 | 39.91 | 47.7 | 2.75 | 0.11 | 4.04 | 34 | 34 | 15 | 3.92 | 3.49 | 3.00 |

**Figure 1.**
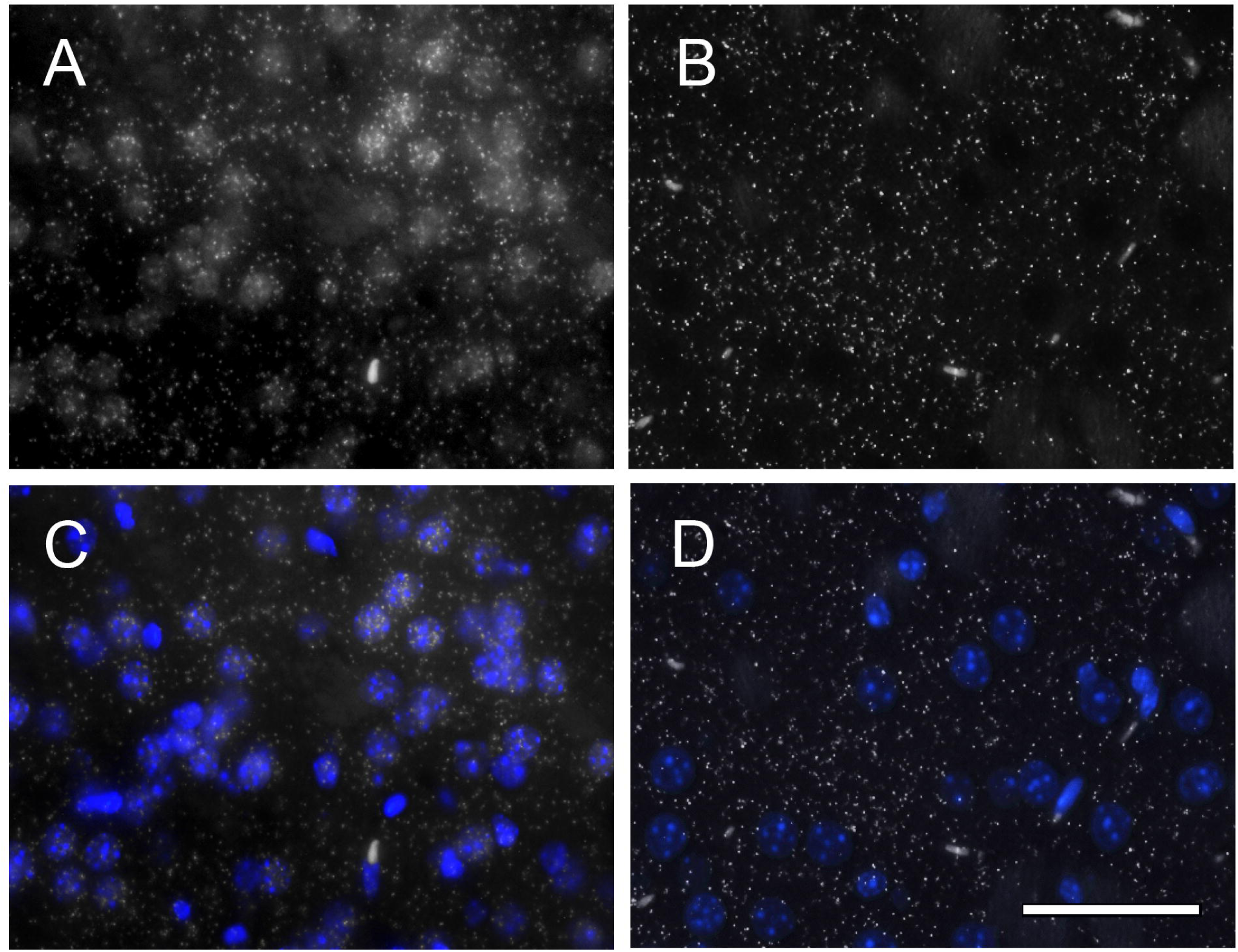
The effect of fixation on the non-specific nuclear signal in single-recognition PLA-FL in mouse brain tissue. Representative images of D2R single-recognition PLA in coronal striatal section from mouse brains fixed by transcardial perfusion of fixative (A, C) or by direct immersion in fixative (B, D). DAPI counterstaining (blue) (C, D) labels nuclei. Note that in the nuclear area, the hazy background signal is almost absent and the puncta overlapping the nuclei are much reduced in the non-perfused tissue compared to the perfused tissue. Scale bar, 50 μm.

### Quantification with Particle Analysis (Image J) and Spot Detector (ICY)

All RGB images were first imported to Image J (*Batch*)[40], transformed to 8-bit images and the contrast was increased 3% (*Enhance Contrast*). To use *Particle Analysis*, an automated (∼170) or manual (110) threshold was applied to all images and then the *Particle size* was set at *1-5 um2*, which was based on manual measurements of PLA puncta from both single and dual PLA assays. The *Overlay* of analyzed particles (PLA puncta) in *ROI manager* was saved and imported to the original images for post-analysis review.

For quantification with ICY (version 1.9.10.0) [41], the RGB images were first transformed to 8-bit images and *Enhance Contrast* (under *Process*) by 3% in Image J. Then PLA puncta were quantified using the *Spot Detector* function in *Detection&Tracking* with the parameters for dual PLA (Detector= scale 1: 334; scale 2 : 60; scale 3 : 100; scale 4 : 1000; Filtering= Min size : 5; Max size : 25) or the parameters for single PLA (detector = scale 1 : 1000 / scale 2 : 89 / scale 3 : 1000 / Scale 4 : 1000; Filtering= Min size : 5; Max size : 25). These parameters were chosen manually on sample images in order to obtain optimal signal-to-noise ratio and then applied to all images for quantification. The names of functions used for image analysis are indicated in italics.

### Quantification with Bayesian Optimized PLA Signal Sorting (BOPSS) package

We designed a custom MATLAB package BOPSS to analyze PLA signal images (https://github.com/neurojojo/BOPSS). BOPSS processes brightfield images containing cells and PLA signal without relying on any user-specified parameters. Instead, clusters of foreground pixels (ROIs) are first selected using an approach typically used in processing histological samples. A subset of these ROIs, chosen at random from both the images of treated and negative control sections, are then used to create twin distributions for 11 intensity-based and morphological features. The remaining ROIs in the dataset are compared against these distributions to assess their likelihood to belong to either category. The two components of BOPSS are further described below:

#### 1. Color-and size-based segmentation of brightfield images into putative nuclei and non-nuclei

Our custom BOPSS package first transforms RGB image pixels from PLA-brightfield images into the LAB color space (CIE International Commission on Illumination 1976). Unlike RGB, which specifies a red, green and blue value for each pixel, the LAB color space uses lightness, as well as green-red and blue-yellow opponency. The transformation between the red-green-blue (RGB) and LAB was accomplished using MATLAB’s built-in *makecform* function. The lightness channel spanned intensities from black (zero) to white (max). Typically, foreground pixels were lighter and had higher values in the lightness channel. Following transformation, a threshold was chosen automatically for each image using MATLAB’s built-in *graythresh* function. Briefly, *graythresh* computes an intensity value which splits the image pixels into two clusters. These two groups are produced to guarantee minimum intraclass variance values as compared to all other possible groupings. The threshold was applied to the lightness channel to select foreground pixels and discard background pixels. The foreground pixels were then selected from the A and B channels. These foreground pixels were next classified as either most likely nuclei or not, based on (1) their values in the color-opponent channels and (2) the size of the clusters they formed in the image. While the first step relied on iterating through k-means results to optimally separate pixels into two groups based on their color, the second step examined the clusters based on their sizes. If a cluster of pixels has a size below the 95th percentile of all cluster sizes over all images, it is deemed a putative punctum. This thresholding based solely on cluster size will tend to produce an upper limit on the number of real puncta as it categorizes any appropriately sized cluster as a punctum. Such clusters generally appear even in negative control images, typically around nuclei. The five percent of clusters with the largest sizes are categorized as nuclei (though they could also be other aggregates) and excluded from further analysis. Undercounting of puncta could occur for images completely lacking nuclei, though the number of puncta excluded from further analysis would be small. Two biological assumptions underlie the performance of this step: first, the majority of puncta should be located outside of nuclei, and second, puncta sizes are far smaller than nuclei. Because there should be virtually no D2R-A2AR dual PLA signal in nuclear areas, we safely exclude the largest pixel clusters in our analyses using BOPSS. By adjusting an internal parameter “Pct_Cutoff”, the user can permit all clusters to pass this first step, regardless of size (this should only be done for images with very few large aggregates of pixels or nuclei). This first step was a highly permissive filter and passed most clusters of pixels that were smaller than nuclei. We referred to these passed clusters as ROI.

#### 2. Naïve Bayesian classifier to categorize PLA puncta and background ROIs

After discarding background and nuclear pixels, BOPSS generates distributions of pixel intensity and shape properties for each of the ROIs. To do this, the program will first label each ROI based on its source image (treated image or a negative control image). BOPSS produces pairs of distributions (treated as well as negative control) over several features of each individual ROI. These features can be broadly described as shape/location features and intensity-based features. The shape/location features calculated for each ROI include: convex area, eccentricity, major axis length, minor axis length, orientation (angle), distance to the nearest neighboring cluster, and density of neighbors. The following intensity-based features are calculated for each ROI: mean pixel value, standard deviation of pixels, entropy of pixels, and range of pixels (highest valued minus lowest valued). Only 50% of the data (training data) is used to generate distributions characterizing the treated and the negative control ROI. In the next step, every remaining ROI (test data) is then classified based on its distance from all of the feature distributions. Each ROI is then classified as either true puncta or noise, depending on whether its features are, on the whole, closer to the center of the distributions for the ROIs extracted from treated images (resulting in true puncta classification) versus ROIs extracted from negative control images (resulting in noise classification). Thus, an ROI’s similarity to the features learned from the training data determine whether it is classified as true puncta or noise.

For the purposes of comparison, we consider “non-optimized” BOPSS (BOPSS_0) puncta to be all ROIs that passed the first step of BOPSS (color- and size-based segmentation), while “optimized” BOPSS puncta were those ROIs that passed both the first and second step (categorization by a naïve Bayesian classifier) (see Results).

### Statistical analysis

Results were analyzed with Graphpad Prism (v5.0.1) and presented as mean ± the standard error of the mean (SEM) or mean± propagation of error, as indicated in figure legends. Two-way ANOVA with post-hoc multiple comparisons test were performed to compare quantification methods.

## Results

### An optimized PLA assay for rodent and human brain sections

Whereas PLA on rodent brain fixed by transcardial perfusion has been widely validated [29, 34], human brain tissue is classically fixed by direct immersion in formalin, which could alter the detection and quality of the signal. Indeed, tissue preparation, especially fixation, is known to alter morphology and immunostaining results [42-44]. To assess how alternative fixation approaches may affect PLA imaging results in human brain sections, we compared the results of immunofluorescent PLA (PLA-FL) on mouse brain sections following different fixation approaches. We performed single-recognition PLA (single PLA), which allows detection of a single antigen with only one primary antibody, for D2R on sections of mouse brains that underwent two different fixation protocols: 1) fixation in 4% PFA overnight after perfusion (perfusion fixation), or 2) fixation by direct immersion in 4% PFA for 4 days without perfusion (direct immersion fixation). With both fixation protocols, we obtained clear PLA signal (puncta) of D2R distributed throughout the striatum, but with a major difference in the non-specific background. As described previously [29, 34, 45] we observed fluorescent signal in the nuclear compartments in samples fixed through perfusion (Fig. 1A, C), whereas this background was much lower in sections from brains fixed without perfusion (Fig. 1B, D). We also performed PLA-FL on sections from snap-frozen mouse brains, which were fixed with 4% PFA after cryosectioning, and observed a similar strong nuclear background (data not shown). These results suggested that the PLA signal for D2Rs from tissue fixed using the direct immersion method is preserved while the nuclear background is greatly reduced. The exact cause of this nuclear background is unclear, but was previously described with PLA-FL from perfused adult animals [29, 46], as well as on brain slices from mouse pups (P0-P1) fixed by direct immersion in fixative [34]. It is therefore tempting to propose that it is the rapid fixation process, which occurs through intracardial perfusion of fixative, or rapid penetration of the fixative in the much smaller pup brain or a thin cryosection, that is responsible for the nuclear background. This signal is unlikely to result from specific or non-specific binding of primary or secondary antibodies, since it is not detectable in regular immunostaining with anti-D2R or anti-A2AR antibodies. Moreover, the nuclear background is still present when antibodies are omitted in the PLA process [29]. We speculate that the crosslinking of chromatin by rapid fixation could result in non-specific binding of the labeled oligonucleotides used during the detection step of the PLA. Regardless of the underlying causes, our results help to explain the dramatic differences in background described in previously published studies [29, 46, 47].

We next performed PLA-FL on paraffin-embedded human brain sections. In marked contrast to mouse brain sections, we observed prominent non-specific signal detectable with both 488 nm and 564 nm lasers (data not shown) in the form of dense aggregates that often accumulated in the soma (Fig. 2A-B), suggesting a non-specific autofluorescence signal, in addition to detection of specific PLA puncta. To address this issue, we replaced the original PLA-FL with a brightfield-compatible detection method, brightfield PLA (PLA-BF), which allows visualization of PLA signal with a chromogenic substrate of horseradish peroxidase (HRP). The PLA-BF for single-recognition of A2AR showed a similar PLA punctate pattern but without the non-specific signal observed with the PLA-FL method (Fig. 2C-D).

**Figure 2.**
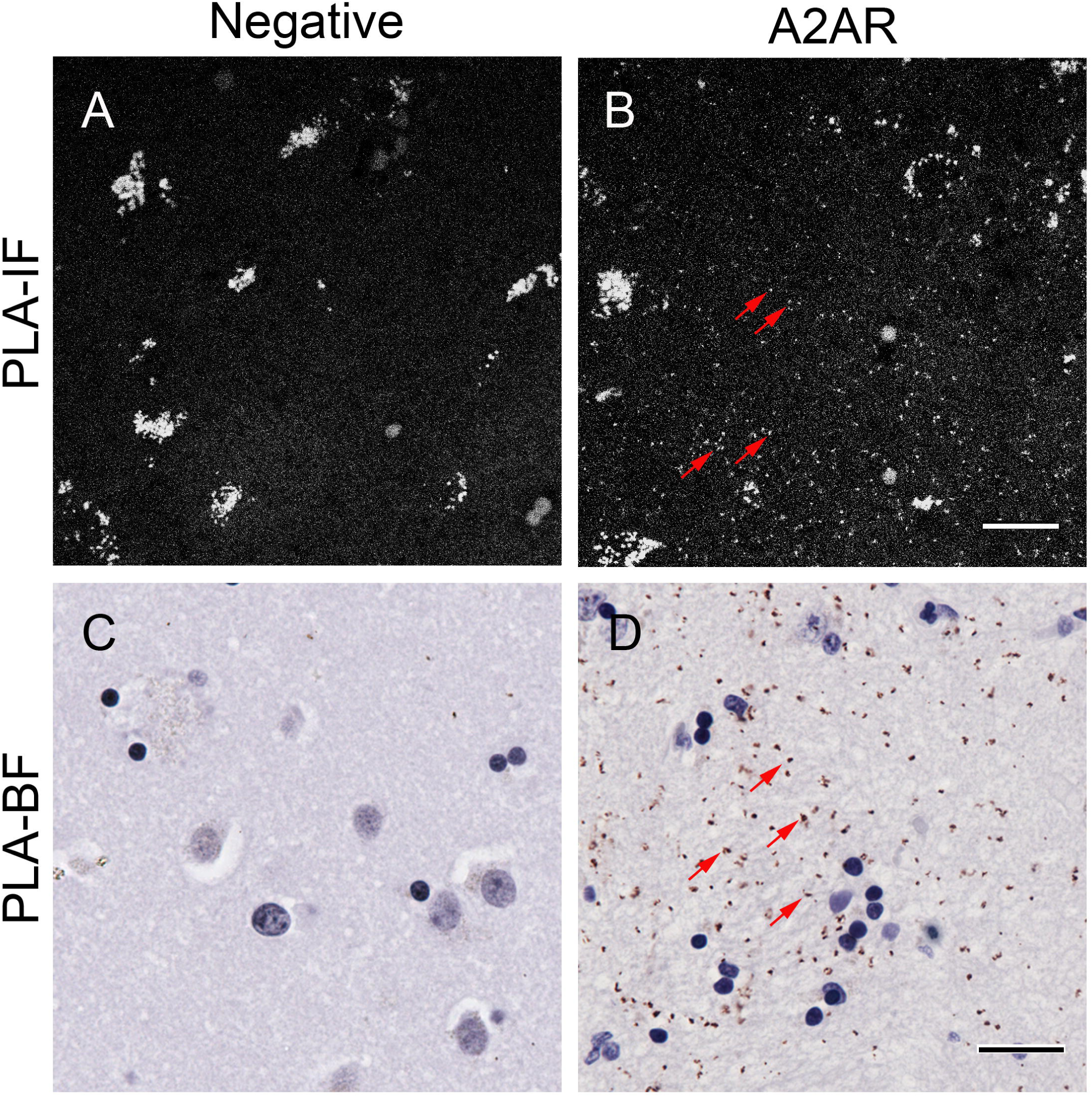
Differences between PLA-FL and PLA-BF detection methods. Representative images of negative controls (A and C) and A2AR single-recognition PLA-FL (B), PLA-BF (D) performed on paraffin-embedded human brain sections of the ventral striatum from the same human subject. The arrows in B and D indicate representative PLA puncta. Hematoxylin counterstaining (blue) labels nuclei (C, D). Scale bar, 20 μm. Note the high autofluorescence background in PLA-FL and the clean background in the PLA-BF.

Autofluorescence is well-known to be a problem in imaging adult human brain but is not an issue with transmitted light, so PLA-BF avoids the autofluorescence that confounds PLA-FL (Fig. 2) and even standard IF in brain samples from human adults and aged animals [48]. Notably, PLA-BF and PLA-FL on mouse brain sections yielded comparable results (data not shown). Altogether, these observations suggest that PLA-BF is a reliable alternative method to PLA-FL in handling paraffin sections from human autopsy brains. Moreover, the stability of brightfield-based signal compared to fluorescence-based methods allows a better conservation of the samples without the photobleaching inherent to the imaging of fluorescent samples.

### Assessing the expression of D2R, A2AR and D2R-A2AR in the human brain

To assess the expression of D2R, A2AR and D2R-A2AR complexes, we performed single PLA-BF for each individual receptor, and dual PLA-BF for D2R-A2AR complexes on coronal sections of adult human ventral striatum that included the nucleus accumbens (NAcc) and limited parts of the putamen (Ptm) and caudate (Cdt). The anatomical structure was confirmed by Luxol fast blue and cresyl violet (LFB/CV) staining, which stains myelin and Nissl substance, respectively (Suppl. Fig. 1A-C). Considering the low number of PLA puncta for D2R-A2AR complexes relative to individual D2R and A2AR in mouse brains [29], we used different concentrations of primary antibodies for dual and single PLAs to obtain optimal staining results while avoiding saturation in single PLA that can lead to overcrowded signal instead of countable puncta. In all tested samples, we observed strong single PLA signal of D2R and A2AR, and relatively weak dual PLA signal of D2R-A2AR complex in the NAcc, Ptm and Cdt (Fig 3 A-I and data not shown). The dual PLA for negative controls that omitted anti-A2AR antibody showed no or few detectable puncta in the same areas (Fig. 3 J-L and data not shown). These results support the existence of D2R-A2AR complexes in native human brains, even though they might be inferred to represent a relatively small fraction of D2Rs and A2ARs.

**Figure 3.**
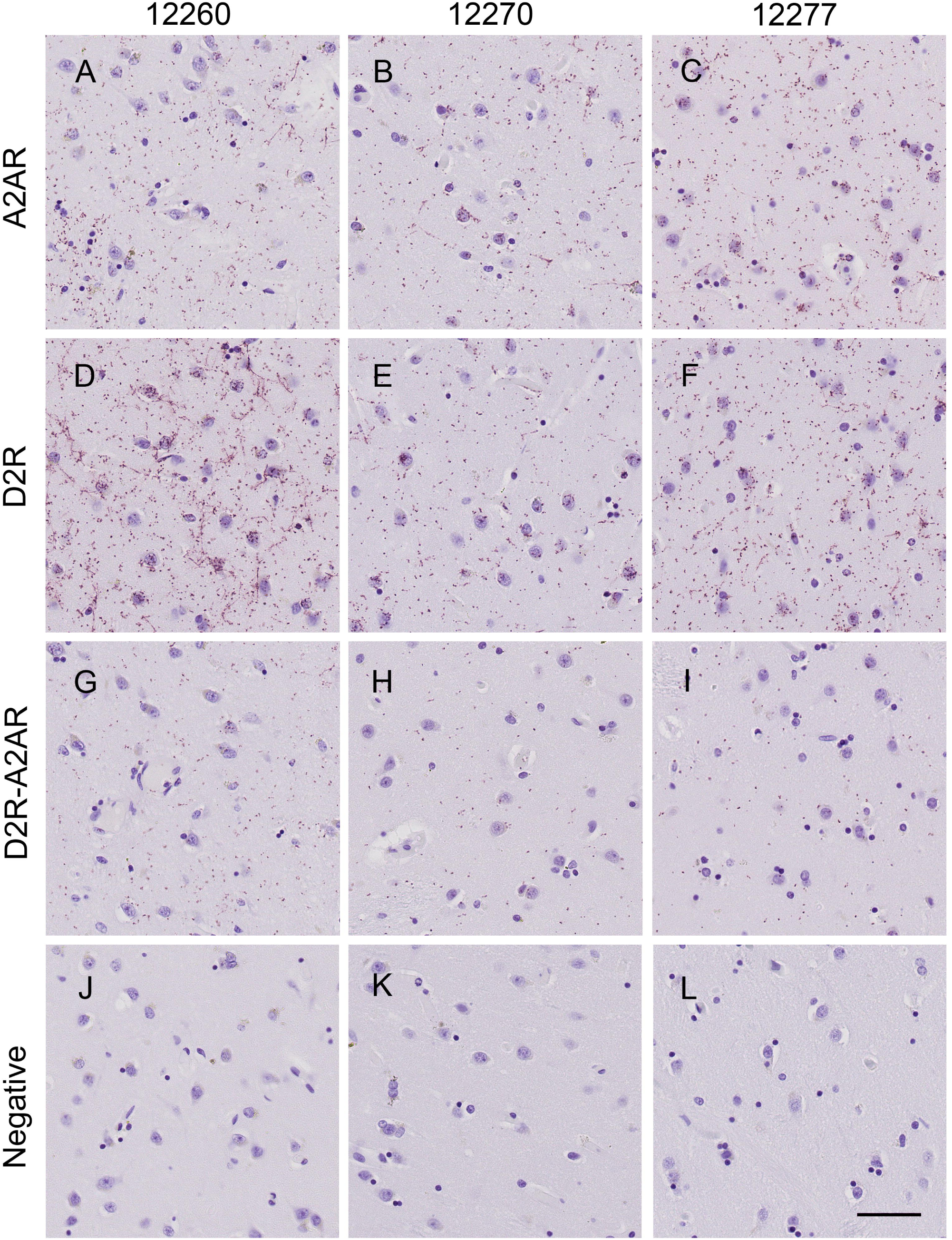
Detection of single PLA for A2AR and D2R, and dual PLA for D2R-A2AR by PLA-BF in the NAcc. The expression of A2AR (A-C), D2R (D-F), D2R-A2AR (G-I) and negative controls (J-L) in the NAcc was detected in three human subjects, PI12260, PI12270 and PI12277. Scale bar, 50 μm.

### Systematic random sampling and automated quantification of PLA-BF signal in human brain sections

Since the PLA signal appears as puncta, quantitative analysis of proteins is more plausible than in traditional immunostaining. However, the size of the human brain tissue makes it challenging to quantify PLA signal from an entire brain region. For example, the brain ROI in this study, the NAcc, was 40.63 ± 0.66 mm^2^ (Table 2). Therefore, we developed a method to allow systematic sampling of a defined brain region. We obtained virtual images by whole slide scanning, divided the ROI into a series of sampling areas, and only quantified PLA signal in the areas (named counting area in this study) that were selected by a systematic random sampling method adapted from Gundersen et al. [39] (Suppl. Fig. 1F). Thus, the total analyzed area represented only 3.36 ± 0.22 % of the ROI (Table 2), but was composed of a set of representative counting areas evenly distributed within the entire ROI region.

To efficiently and reproducibly quantify PLA puncta across our large dataset in an unbiased way, we applied and compared three different quantification approaches. Two of these approaches, *Analyze Particles* (ImageJ) [40] and *Spot detector* (ICY) [41], rely on the user to set intensity and size parameters to distinguish clusters of pixels against the background of an image. In contrast, our custom MATLAB package BOPSS that we designed to automatically quantify PLA-BF signal is an unsupervised machine learning approach and requires no parameters to be set by the user.

∘ *Analyze Particles* (Image J) depended on *Threshold* and *Particle Size* to detect PLA puncta. The automated threshold (Fig. 4: ImageJ_auto) worked well for dual PLA signal, but underestimated the signal for single PLA, and showed high counts in the negative controls (Fig. 4D-F, P), which resulted in low signal (the dual PLA signal)/background noise (the negative control signal) (S/N=2.14). Most of the non-specific puncta in the negative controls detected by *Analyze Particles* corresponded to background in the nuclear area, typical of the signal obtained with classical DNA intercalating reagents like DAPI or Hoechst. Manually optimizing the threshold reduced the non-specific detection in the negative controls, but caused more severe underestimation for both dual and single PLA, and failed to improve S/N (data not shown). Adjusting the size of particles also failed to improve S/N or underestimation of specific PLA signal (data not shown). These results suggested that, *Analyze Particles* (Image J) cannot properly discriminate specific and non-specific signals and underestimated specific PLA signal.

**Figure 4.**
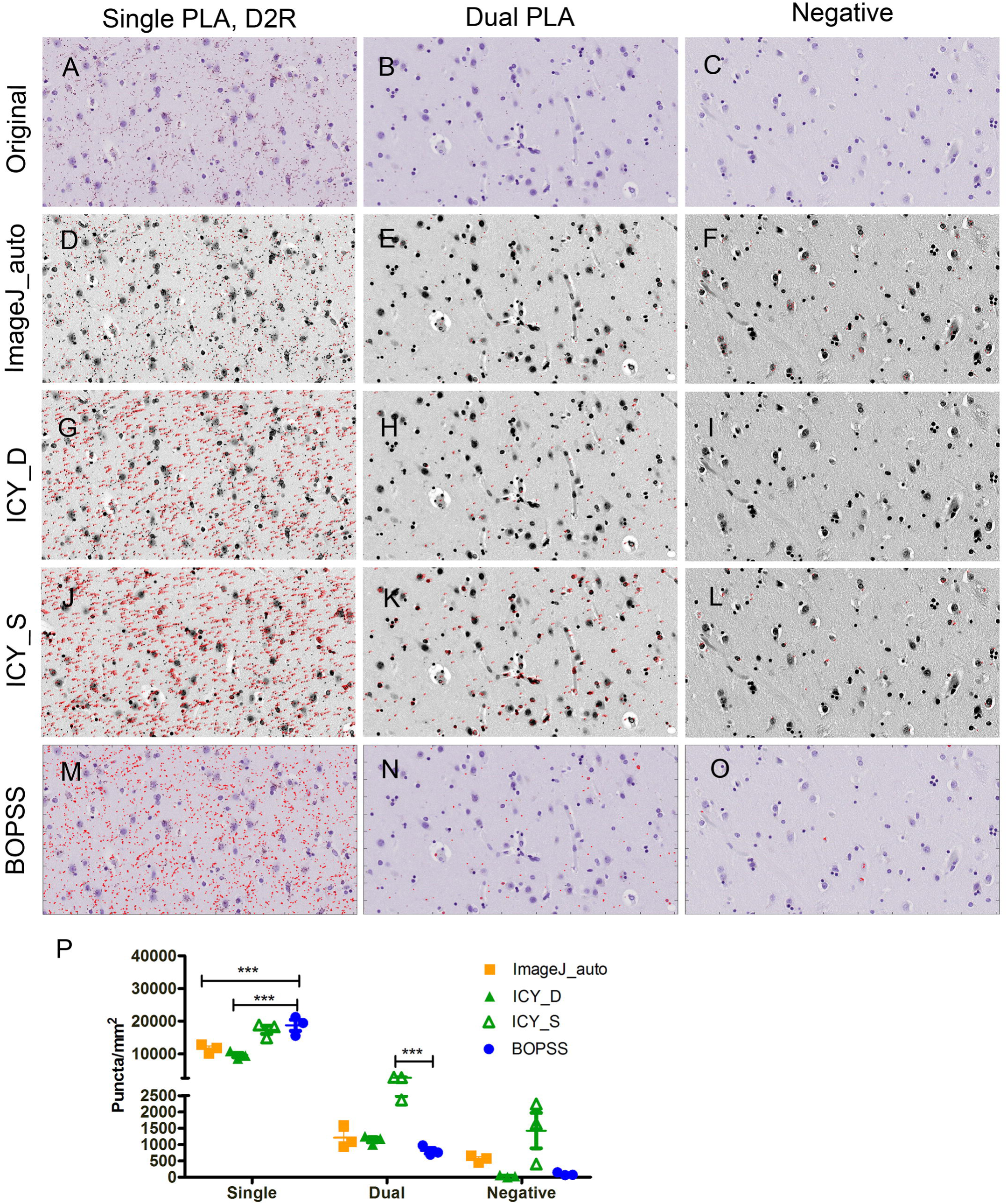
Comparison of automated quantification methods. Three counting images of the dual (A), negative (B) and single PLA (C) from subject PI12277 were used to test puncta detection and quantification approaches with selected parameters: *Particle Analysis* (Image J) with auto threshold (Image J_Auto, D-F), *Spot Detector* (ICY) with parameters favoring detection of either dual and negative (ICY_D, G-I) or single (ICY_S, J-L) PLA signals, and BOPSS (BOPSS, M-O). Image J and ICY quantified the puncta in the transformed and contrast enhanced images (D-L) as described in the methods. The counted puncta were marked in red dots (D-F, M-O) or labeled in red numbers (G-L) in the analyzed images. Two-way ANOVA and Bonferroni post-tests were performed to compare the results of BOPSS to those of the other methods (P). n=3,* P value <0.05,*** P values <0.01. Data were plotted as mean ± SEM.

∘ *Spot detector* (ICY) detected PLA puncta through a series of functions: 1) *Scales* that determine the sensitivity and the size of spots to detect, and 2) a *Filtering* that determines the maximum and minimum size of accepted objects. When parameters were optimized for dual PLA, this method showed very little – if any-non-specific detection in the negative controls and good detection for dual PLA, but noticeable underestimation for single PLA (Fig. 4G-I, P: ICY_D). With optimized parameters for single PLA, *Spot Detector* showed increased detection in every PLA assay, including non-specific detection in the negative controls (Fig. 4J-L, P: ICY_S). Therefore, two different sets of parameters were necessary for specific and efficient detection for single PLA (puncta of high density) and dual/negative PLA signal (puncta of low density).

∘ In contrast to ImageJ and ICY, BOPSS uses a machine learning algorithm to learn the properties of the puncta based on the data. It measures a variety of features for each cluster of pixels and uses the distributions of these features to contrast between negative controls and samples. It processes color images containing cells and PLA signal without relying on any user-defined parameters, but rather on the specification of negative control and experimental images. On a standard laptop PC (2.6 GHz CPU), we analyzed output results for ten megabytes of images in five seconds. In a pretest with 3 full counting images of each PLA condition, single PLA, dual PLA and negative control, this all-in-one script completed machine learning based optimization and quantification automatically. Compared with the pre-optimization counting (BOPSS_0) that only selected puncta based on size and color, the optimized quantification BOPSS (BOPSS) improved the signal (the dual PLA signal)/background noise (the negative control signal) ratio of dual PLA from 1.63 to 8.41, reduced the averaged counts per image in the negative controls from 129± 33 to 10± 3, which represents a 91% reduction in false positives, and maintained efficient detection in both dual and single PLA (Suppl. Fig. 2M). To assess the accuracy of BOPSS, we randomly selected three areas (Suppl. Fig. 2 A-C), which covered 43 % of a full counting image, and counted the PLA puncta manually (Suppl. Fig. 2 J-L) or with BOPSS (BOPSS_0 and BOPSS, Suppl. Fig. 2 D-I). The results showed that BOPSS achieved remarkably similar results for dual PLA and negative control compared with the manual counting (Suppl. Fig. 2N). For single PLA, BOPSS counted slightly more PLA puncta than the mean of three independent manual counting attempts (Suppl. Fig. 2N). However, manual counting, while highly reproducible for the low density signals, was quite variable and extremely difficult for the high density puncta. Post hoc manual analysis verified that although BOPSS occasionally undercounts PLA puncta for high density signals (Suppl. Fig. 2G), it also tended to occasionally overcount signal in nuclear areas (Suppl. Fig. 2G-I).

Compared with the other two automated quantification methods described above, BOPSS avoided time-consuming and potentially biased parameter optimization, but nonetheless resulted in comparable quantification results for the dual PLA and negative controls as Image J_auto and ICY_D (with optimized parameters for single PLA) (Fig. 4P). At the same time, BOPSS significantly improved detection of dense single PLA signal, leading to comparable results for single PLA as ICY_S (with optimized parameters for single PLA) (Fig. 4M-P, BOPSS). Therefore, BOPSS achieved a favorable S/N for dual PLA by reducing non-specific detection without underestimating real PLA puncta, and simultaneously overcame the underestimation issue for single PLA. In addition, BOPSS led to comparable results as manual counting of PLA puncta (Suppl. Fig. 2), and was much more consistent than manual quantification, which is an advantage for processing a large dataset in an unbiased manner. Thus, overall, BOPSS seems to be a straightforward approach for achieving completely automated, efficient and specific detection of PLA signals and worked equally well for both single PLA (puncta of high density) and dual (or negative) PLA signal (puncta of low density).

PLA has been previously applied to postmortem human brains in a very limited number of studies [35, 36] and optimization of unbiased quantification methods has not been achieved. Combining whole slide scanning microscopy, the SRS sampling method, and automated quantification with BOPSS, we developed here a systematic automated approach for quantitative study of PLA-BF, which enhanced our data analysis capacity to the level required for processing image data from the human brain.

### Quantifying the expression of D2R, A2AR and D2R-A2AR in the human brain

Because the ventral striatum tissue blocks contained only partial Ptm and Cdt of varied sizes, we focused our quantification in the NAcc. The quantification results from the three human subjects (Table 1) showed varied counts of D2R, A2AR and D2R-A2AR complex in the NAcc (Fig. 5 A-C). The averaged relative fractions of D2R-A2AR complex were 12.3±1.8 % relative to total A2AR and 13.4±5.3 % relative to total D2R. In this small preliminary analysis, we are unable to assess whether the differences in the absolute levels of complex and their levels relative to the two single receptors represent true individual variation or subtle differences in the exact regions quantified (Fig. 5 E, F). Nonetheless, the results show that our method can quantify both absolute numbers of antigens as well as the proximity between the two antigens.

**Figure 5.**
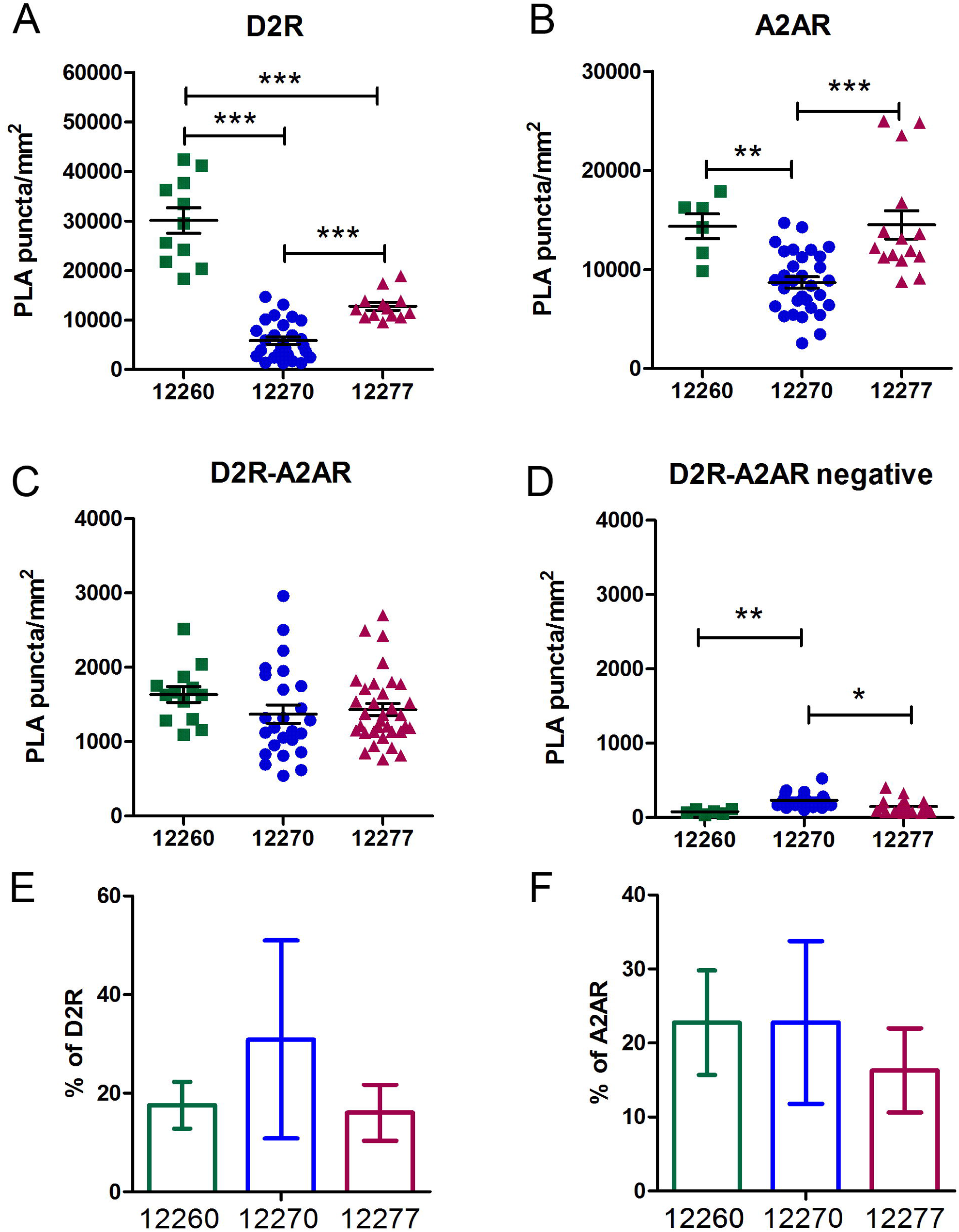
Quantifying the signal of single PLA for A2AR and D2R, and dual PLA for D2R-A2AR, in the NAcc. The numbers of PLA puncta / mm^2^ were quantified by BOPSS (A-D, data were plotted as mean ± SEM). The fractions of D2R-A2AR dual PLA puncta relative to A2AR (E) and to D2R (E) single PLA were calculated with the means in (A-C) and data were plotted as mean ± propagation of error.

The existence of heteromers composed of A2AR and D2R has been supported by multiple studies using different approaches, including co-immunoprecipitation *in vitro*, FRET and BRET in living cells, and PLA-FL in rodent striatum, making it the most characterized GPCR heteromer [16-19]. In this study, we have, for the first time, detected D2R-A2AR complexes *in situ* in postmortem human brains, suggesting that the physical proximity between D2R and A2AR is conserved from rodent to human. Whether this signal represents true heteromerization of two functional receptors that communicate directly or close proximity of receptors in microdomains or a larger complex remains to be determined. We have explored systematic sampling and automated quantification methods to assess the expression of D2R, A2AR and D2R-A2AR among individuals. While the number of samples here was very small, the results demonstrate that single and dual PLA assays can be carried out in paraffin sections of human autopsy brains, that the signal can be measured by computerized image analysis, and that the results can be used reliably to explore differences in protein expression in a way that is vastly more quantitative than traditional immunostaining.

## Acknowledgements

This study was supported by MH54137 (Javitch), MH060877 (Dwork) and MH090964 (Mann). We thank Dr. Gorazd Rosoklija and staff at the Core Facility, Department of Pathology, HICCC, Columbia University Medical Center. for technique support.

