## Supplementary material for "Detecting GPCR Complexes in Postmortem Human Brain with Proximity Ligation Assay and A Bayesian Classifier": Suppl.

`

**Supplemental Methods and Materials**

*Systematic Randomly Sampling*

The systematic random sampling for human brain PLA quantification was performed according to the following protocol.

1. Outline the territory of interest for each sample. Open a virtual whole slide image (scn file) of LFB/CV staining in Leica SCN400 Image Viewer (SCN viewer, version 2.2), zoom to have a full view image that fits the screen, and export this full view image as a tiff file with the ROI image export function. Open this full view image in Photoshop (Adobe), create a new layer, draw outlines of brain section and sub-territories on the separate layer, and print the outline layer on transparency film without scaling (Suppl. Fig.3D).
2. Make a sampling grid. In Adobe Illustrator, create a grid (16.5 mm x 16.5 mm/cell in this protocol) image of the same size as the outline image in step (1). Print the grid on transparency film without scaling. This printout is the fractionator for sampling (Suppl. Fig. 1E), which can divide an ROI into a series of sampling areas. Each of the grid cells corresponds to one sampling area. The size of the grid cell must be adjusted to divide the ROI into a suitable number of sampling areas.
3. Select and mark the counting locus. Overlap the films of outline (step 1) and the sampling grid (step 2). The brain section or region of interest (ROI) is thus divided into a number of sampling areas. Mark the up-left corner of each cell as the counting locus (Suppl. Fig. 1E), which can be anywhere inside a sampling area, but need to be consistent for all sampling area.
4. Sampling. Open a whole slide image (scn file) of PLA in SCN Viewer, and zoom it to get a full view image (0.4x in this study) that fits your screen. Overlap and mount its outline film from step (1) on screen. Mount the film of the sampling grid (step 2) on top of the outline film to cover the entire ROI. Place the mouse indicator at one counting locus, which is marked in step3, scroll the middle button of the mouse to zoon the image to 40X, and right click to export ROI image at 40x as a tiff file. This is a full magnified counting image and will be analyzed as representative of the sampling area. Repeat this for all sampling areas inside the ROI. A counting area is only analyzed when it locates inside the ROI.
5. Examine the quality of the images and exclude unfocused or oversaturated images because they will not be analyzed correctly.

**Supplemental Figure Legends**

**Suppl. Figure 1.** Sampling procedures for PLA-BF. Luxol fast blue/cresyl violet staining was performed to show the white matter and the grey matter (A-C). An outline of ventral striatum sub-territories (D) was drawn based on the luxol fast blue/cresyl violet staining for each sample. The overlap of the fractionator (grid) (E) and the outline (D) divided the region of interest (ROI) into a number of evenly distributed sampling areas (F). One counting area was selected in each sampling area (indicated by *) and the 40x image of this counting area was exported for quantification. The sampling area was only counted when its counting area was inside the ROI.  Scale bar, 5 mm.

**Suppl. Figure 2.** Quantify PLA signal through BOPSS and manual counting. Three randomly selected areas in a full counting image of each PLA conditions, single (A), dual (B) and negative PLA (C) (from PI12277) were quantified with BOPSS or manually (three times independently).  The puncta counted by BOPSS were marked in red in the representative images of pre-optimization (BOPSS_0, D-F) and post-optimization analysis (BOPSS, G-I). The blue arrows indicated the examples of reduced non-specific detection in post-optimization analysis. The manually counted puncta were marked in black and labelled with yellow numbers with Cell Counter (Image J) (J-L). The red and white arrows indicated examples of over-counting and under-counting detection by BOPSS compared with manual counting, respectively.

Suppl. Fig. 1.

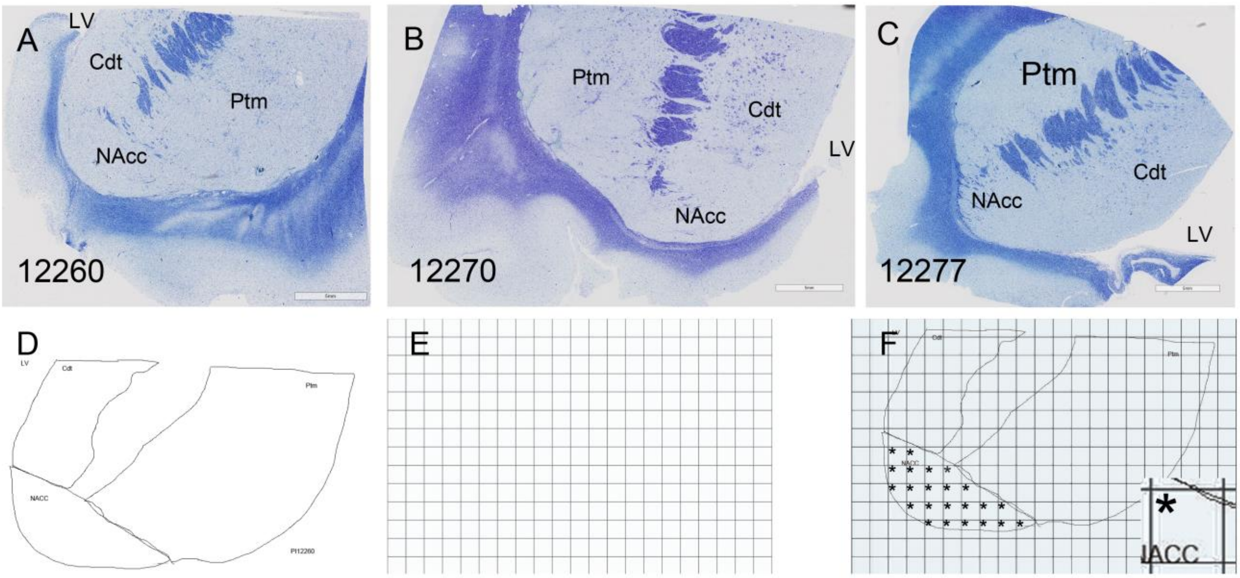
A
LV
Cdt
Ptm
NAcc
12260
B
Ptm
Cdt
LV
NAcc
12270
C
Ptm
Cdt
NAcc
LV
12277
D
LV
Cdt
Ptm
NACC
PI12260
E
F
Cdt
Ptm
NACC
ACC

Suppl. Fig. 2

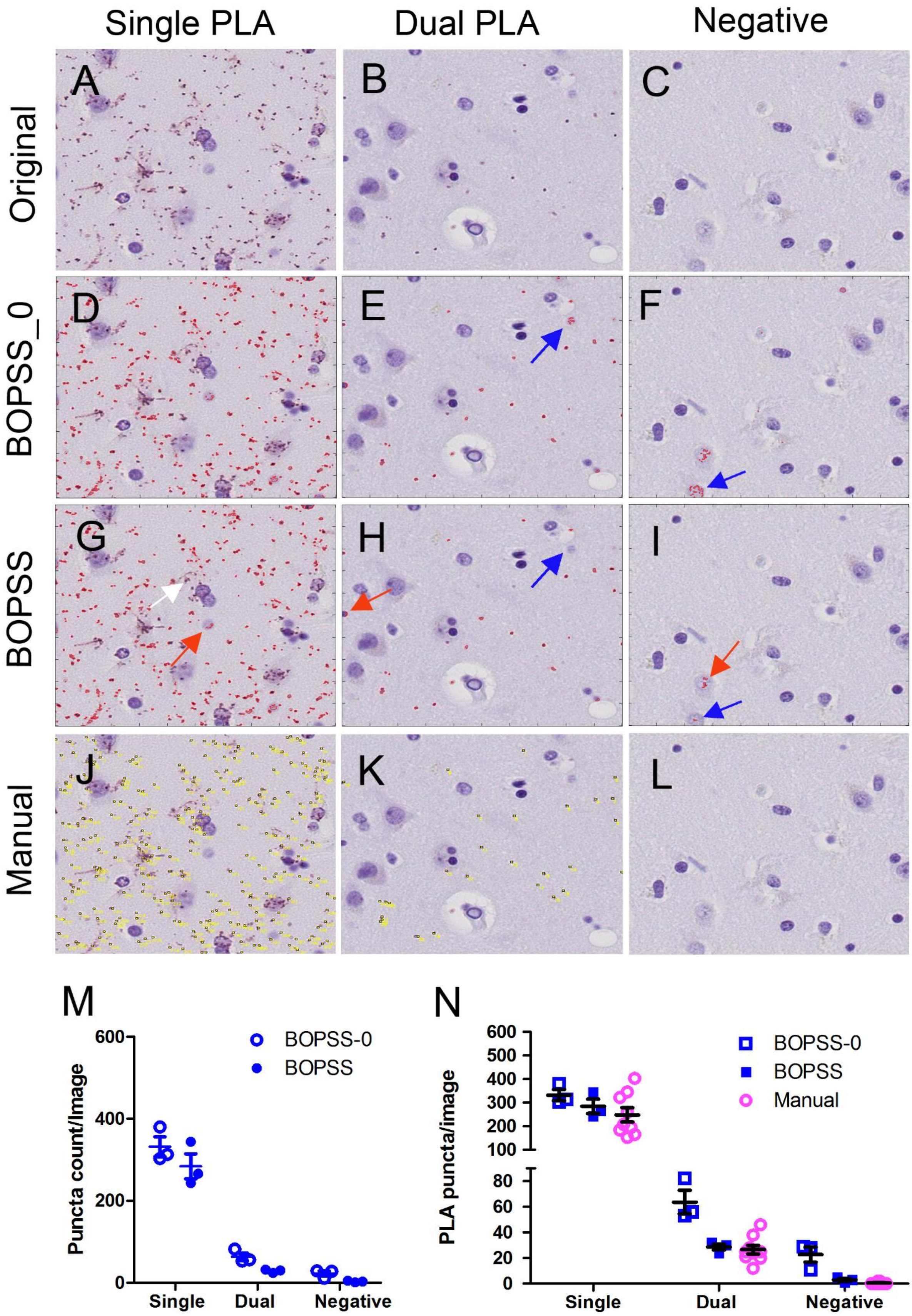

Single PLA
Dual PLA
Negative
Original
BOPSS_0
BOPSS
Manual
A
B
C
D
E
F
G
H
I
J
K
L
M
N
Puncta count/Image
600
400
200
0
BOPSS-0
BOPSS
Single
Dual
Negative
PLA puncta/image
600
500
400
300
200
100
80
60
40
20
0
BOPSS-0
BOPSS
Manual
Single
Dual
Negative
